# Acute induction of anomalous blood clotting by molecular amplification of highly substoichiometric levels of bacterial lipopolysaccharide (LPS)

**DOI:** 10.1101/053538

**Authors:** Etheresia Pretorius, Sthembile Mbotwa, Janette Bester, Christopher Robinson, Douglas B. Kell

## Abstract

It is well known that a variety of inflammatory diseases are accompanied by hypercoagulability, and a number of more-or-less longer-term signalling pathways have been shown to be involved. In recent work, we have suggested a direct and primary role for bacterial lipopolysaccharide in this hypercoagulability, but it seems never to have been tested directly. Here we show that the addition of tiny concentrations (0.2 ng.L^−1^) of bacterial lipopolysaccharide (LPS) to both whole blood and platelet-poor plasma of normal, healthy donors leads to marked changes in the nature of the fibrin fibres so formed, as observed by ultrastructural and fluorescence microscopy (the latter implying that the fibrin is actually in an amyloid β-sheet-rich form. They resemble those seen in a number of inflammatory (and also amyloid) diseases, consistent with an involvement of LPS in their aetiology. These changes are mirrored by changes in their viscoelastic properties as measured by thromboelastography. Since the terminal stages of coagulation involve the polymerisation of fibrinogen into fibrin fibres, we tested whether LPS would bind to fibrinogen directly. We demonstrated this using isothermal calorimetry. Finally, we show that these changes in fibre structure are mirrored when the experiment is done simply with purified fibrinogen and thrombin (± 0.2 ng.L^−1^ LPS). This ratio of concentrations of LPS:fibrinogen *in vivo* represents a molecular amplification by the LPS of more than 10^8^-fold, a number that is probably unparalleled in biology. The observation of a direct effect of such highly substoichiometric amounts of LPS on both fibrinogen and coagulation can account for the role of very small numbers of dormant bacteria in disease progression, and opens up this process to further mechanistic analysis and possible treatment.

**Significance statement:** Most chronic diseases (including those classified as cardiovascular, neurodegenerative, or autoimmune) are accompanied by long-term inflammation. Although typically mediated by ‘inflammatory’ cytokines, the origin of this inflammation is unclear. We have suggested that one explanation is a dormant microbiome that can shed the highly inflammatory lipopolysaccharide LPS. Such inflammatory diseases are also accompanied by a hypercoagulable phenotype. We here show <u>directly</u> (using 6 different methods) that very low concentrations of LPS can affect the terminal stages of the coagulation properties of blood and plasma significantly, and that this may be mediated via a direct binding of LPS to a small fraction of fibrinogen monomers as assessed biophysically. Such amplification methods may be of more general significance.

## INTRODUCTION

‘LPS’ describes a variety of cell wall lipopolysaccharides shed by Gram-negative bacteria; also known as ‘endotoxin’, they have been found in various fluids, including whole blood. The ‘concentrations’ are typically assayed using the *Limulus* amoebocyte lysate assay (e.g. [1–3]). However, although satisfactory in simple matrices, this test is not considered very reliable in blood [4, 5]. Indeed, because it is so hydrophobic, little or no LPS is actually free (unbound), and so it is not even obvious what its ‘concentration’ in blood might mean (see [5]). Although the quantitative assessment of LPS concentrations in whole blood can be problematic, its presence in this fluid may have important and clinically relevant effects on the blood microenvironment, and may be central in the treatment of inflammatory conditions [5–8].

LPS molecules are potent inflammagens (e.g. [9–11]) and may be both cytotoxic and/or neurotoxic [5, 12–15]. They are known to induce the production of a variety of pro-inflammatory cytokines [16–19] that are involved in various apoptotic, programmed necrosis and pyroptotic pathways [5, 20, 21]. Indeed, cytokine production [16] is central to the development of inflammation [22]. A further characteristic of systemic inflammation is a hypercoagulatory state [23–27]. Such hypercoagulability is a common pathology underlying all thrombotic conditions, including ischemic heart disease, ischemic stroke, and venous thromboembolism [28]. Furthermore, a hypercoagulable state is typically associated with pathological changes in the concentrations of fibrin(ogen) [29, 30], and in particular an increase in the level of the fibrin degradation product D-dimer is seen as a reliable biomarker for cardiovascular risk [31, 32].

As well as its cytokine-dependent effects, the question then arises as to whether LPS can cause hypercoagulation by acting on the coagulation pathway more directly. One route is via tissue factor (TF) upregulation; TF is related to the cytokine receptor class II family, and is <u>active early</u> in the (extrinsic) coagulation cascade, where it is necessary to initiate thrombin formation from prothrombin [33]. Recently, it was shown that LPS may upregulate TF; 100 ng.mL^−1^ LPS added to healthy cord whole blood of newborns, or the whole blood (WB) of healthy adults, induced tissue factor-mediated activation of hemostasis [34]. LPS from *Escherichia coli* (100 ng.mL^−1^) also activated the coagulation system when added to WB, via a complement-and CD14-dependent up-regulation of TF, leading to prothrombin activation and hypercoagulation [35]; however, this was noted after 2 hours, and therefore it was not an acute process [35]. Note that in these studies, the anticoagulant was lepirudin which prevents thrombin activation such that the effects of thrombin could not be evaluated. In the present work we used citrate as an anticoagulant.

It occurred to us that, in addition to changes in TF expression by LPS, the process might also involve the direct binding of the lipophilic LPS to circulating plasma proteins, particularly fibrinogen, and that this (potentially rapid) binding might also cause pathological changes in the coagulation process. This would be independent of the slower TF activation, and thus an acute and relatively immediate process (Fig 1). This indeed turned out to be the case.

**Fig 1:**
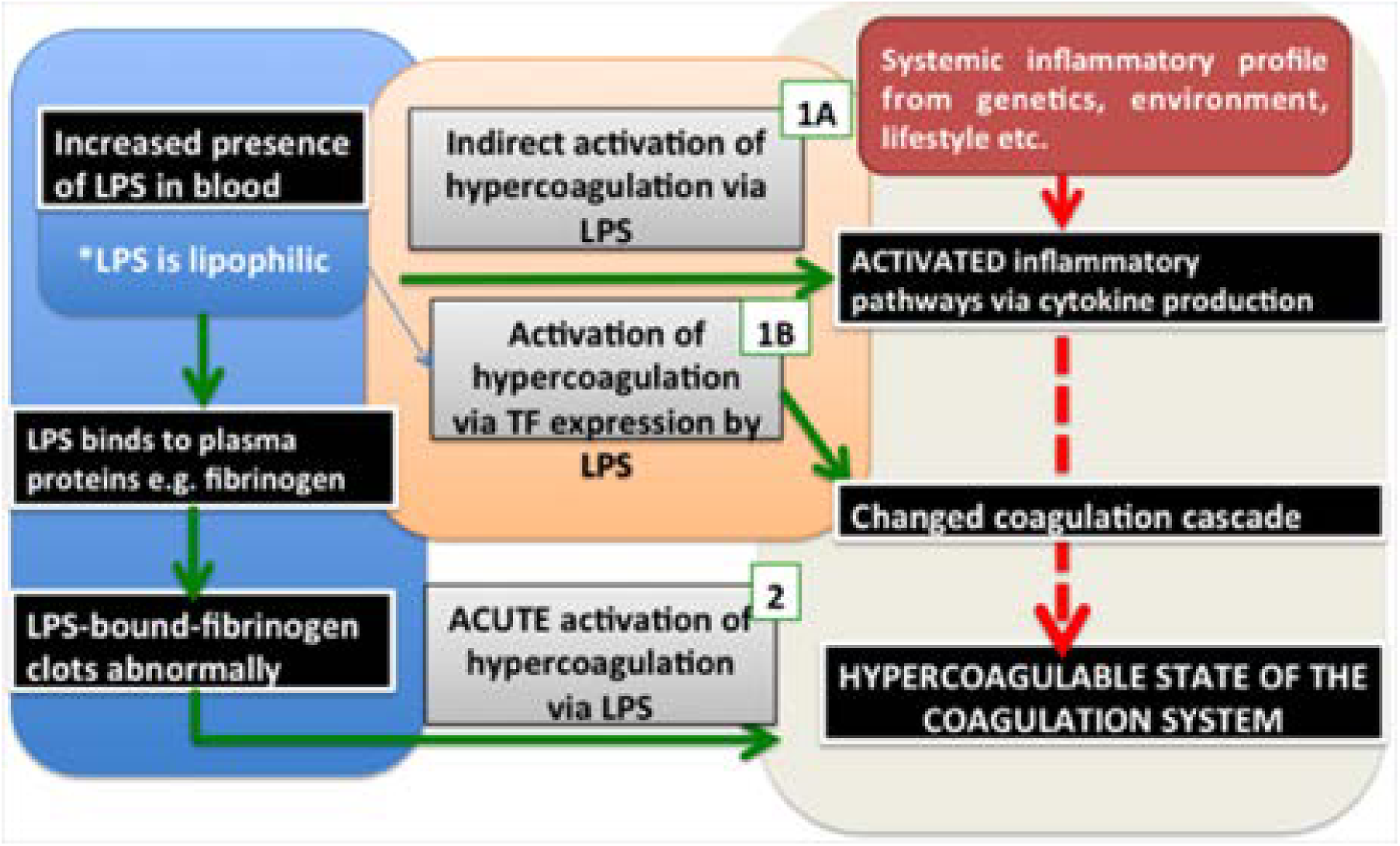
High-level effects of systemic inflammation on the coagulation system and the pathologic effects of LPS when present in blood, and how it influences coagulation via a direct or indirect activation. **Processes 1A and B** are currently known for LPS activity in blood, while **process 2** is a newly proposed acute reaction effect of LPS on blood and plasma.

## RESULTS

### Scanning electron microscopy of whole blood, plasma and purified fibrinogen

To investigate our hypothesis that LPS may cause hypercoagulation via an acute, and direct binding reaction (by interaction with plasma proteins directly involved in the clotting cascade), we investigated the effect of 2 LPS preparations from *E. coli* (viz. O111:B4 and O26:B6). These were added to whole blood (WB) of healthy individuals, to platelet (and cell) poor plasma (PPP), and to purified fibrinogen.

Although the physiological levels of LPS are said to be 10 - 15 ng.L^−1^, and little or none of it is free [5], in our hands the addition of LPS at these concentrations caused immediate coagulation when they were added to whole blood. Fig 2 shows the effect of 0.2 ng.L^−1^ O111:B4 LPS when added to WB and incubated for 10 min. Dense matted deposits are spontaneously formed; these are not seen in healthy whole blood. Fibrinogen with added O111:B4 or O26:B6 LPS with just 30s exposure (no thrombin added) also spontaneously formed matted deposits (results not shown).

**Fig 2:**
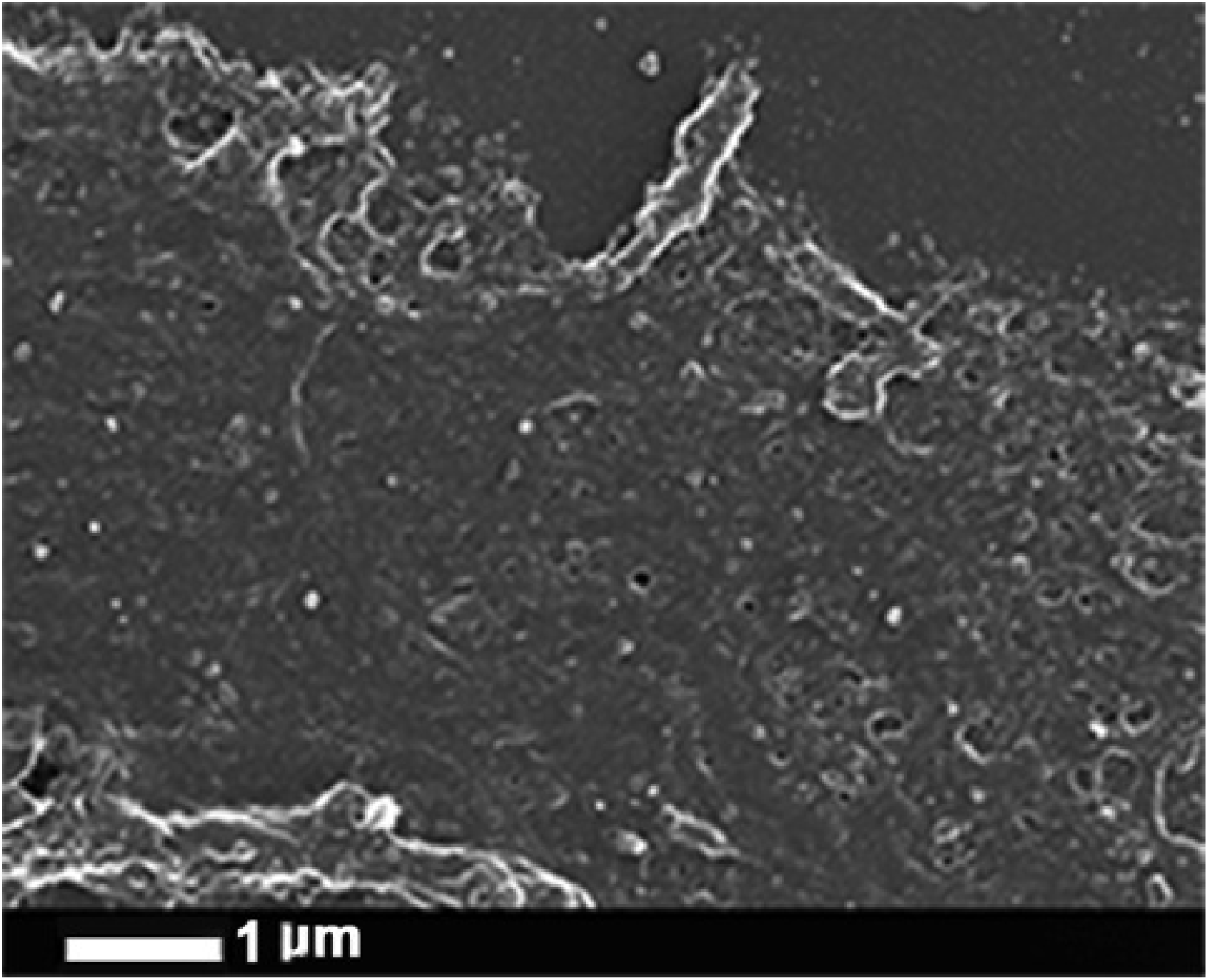
Effect of O111:B4 LPS (0.2 ng.L^−1^) on whole blood (without thrombin), where dense matted deposits were spontaneously formed, not seen in control whole blood smears.

### PPP and LPS

Fig 3 shows the effects of thrombin on clot formation for a healthy control (3A) and when the PPP was pre-incubated for 10 min with 0.2 ng.L^−1^ O111:B4 LPS (two examples in 3B, C). Fig 3D shows the distribution of fibre thicknesses for the 30 individuals, with and without added LPS. The fibre thickness is much more heterogeneous after LPS is added; Clearly these tiny amounts of LPS are having enormous effects on the clotting process. These kinds of netted structures, that we have also termed ‘dense matted deposits’, were previously seen in inflammatory conditions such as diabetes [36–38], iron overload and stroke [36, 39–41]. Typically healthy fibrin fibre networks form a net where individual fibrin fibres are seen, but with added LPS a matted net develops. There is a significant difference (p < 0.0001) between fibre thickness before and after LPS treatment in the presence of thrombin. Note, though, that the distribution of the fibre thickness in the LPS-treated group varies from very thick to very thin. In some cases, continuous fibre plates are formed, where no individual fibres could be seen or measured. This explains the difference in n between the ‘before’ and ‘after’ treatment (1450 versus 1330 measured fibres).

**Fig 3:**
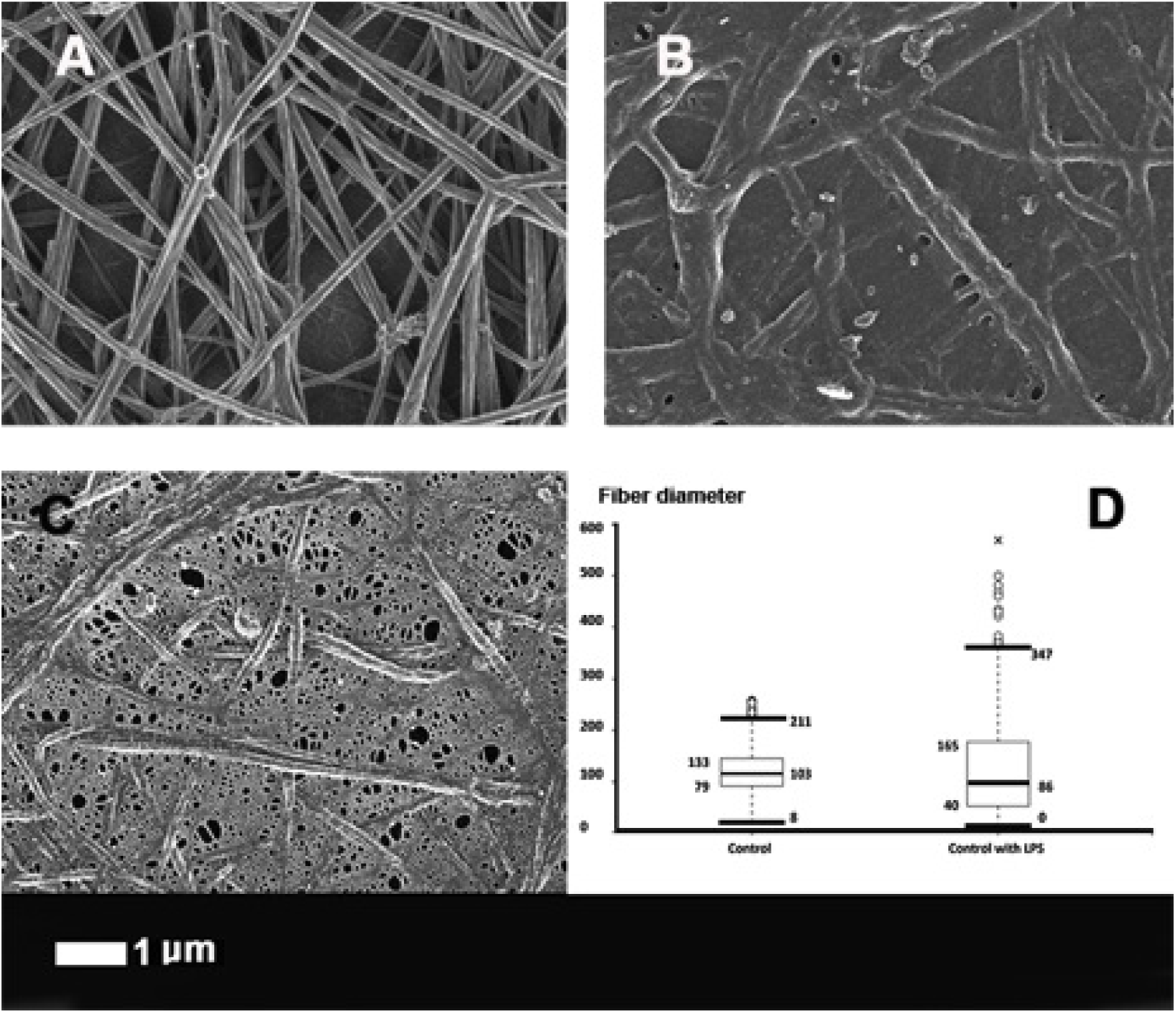
The effect of 0.2 ng.L^−1^ O111:B4 LPS on the morphology of fibrin fibres in the platelet poor plasma (PPP) of healthy individuals (with added thrombin). **A)** Healthy fibres; **B and C)** PPP with added LPS. **D)** Fibre distribution of the control fibres and of controls with added LPS of 30 individuals. Note: in samples with added LPS there were areas of matted layers with no visible fibres to measure. Fibres were measured using ImageJ as described in Materials and Methods.

The experiment with the O111:B4 LPS was also repeated with a shorter 30s exposure time. PPP with LPS and added thrombin showed fibre agglutination starting to happen in only 30s of LPS exposure. These shorter experiments are to be contrasted with previously reported experiments that showed the much longer-term effect of LPS, involving cytokine production, including increased TF production via monocytes. By adding LPS to PPP (with thrombin) we bypass the possibility that LPS can stimulate TF production via the monocyte route suggested by [35, 42]. To determine if another type of LPS would also cause the changes noted above, we also added *E. coli* O26:B6 LPS to PPP of 5 individuals (30s and 10 min exposure time), followed by addition of thrombin. SEM results showed the same trends as noted with the O111:B4 LPS (results not shown).

### Confocal microscopy

The reaction, and presumed binding, of the hydrophobic LPS within fibrinogen fibres implies that they contain, or expose, significant hydrophobic elements. Such elements, also common in amyloid-like fibrils [43], can be stained fluorescently using dyes such as thioflavin T (TFT) [44]. We therefore studied the effect of 0.2 ng.L^−1^ LPS on the ability of the fibrin fibres formed following thrombin treatment of PPP to bind TFT (Fig 4). In contrast to the LPS-free controls (4A) there is a very substantial binding of TFT to the fibrin fibres formed in the presence of the LPS (4B).

**Fig 4:**
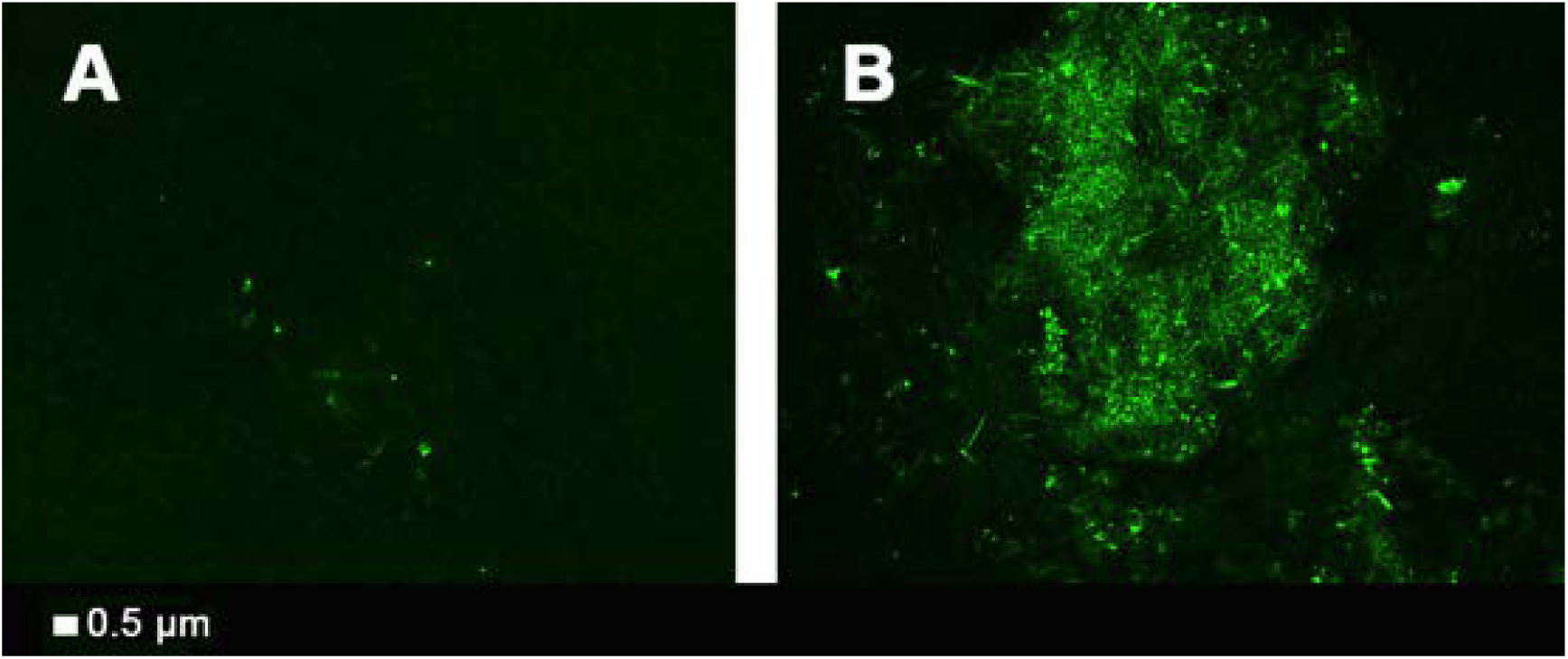
**A)** Control PPP with TFT and thrombin and **B)** as A but pre-incubated with 0.2 ng.L^−1^ LPS.

### Purified fibrinogen

It is worth rehearsing just how big an effect this is in molar terms: fibrinogen (MW 340 kDa) is present at in plasma at concentrations of ~2-4 g.L^−1^ (Weisel’s authoritative review [45] gives 2.5 g.L^−1^), and its levels are increased during inflammation (see above), while the LPS (MW 10-20kDa) was added at a concentration of 0.2 ng.L^−1^. We will assume 15kDa for the MW of LPS and 30 fg LPS.cell^−1^. Thus 0.2ng.L^−1^ = 13 fM and 2.5 g.L^−1^ fibrinogen ~ 7.35 μM which is a molar ratio of LPS;fibrinogen monomer in the whole blood of less than 10^−8^:1. Since we are here only looking at the terminal stages of clotting, we considered that fibrinogen might be an important mediator of the LPS-induced hypercoagulation. Thus, we also added both LPS types to purified fibrinogen (30s and 10 min exposure time) with added thrombin. Even the 30s exposure time changed the fibrin fibres to form fibrils or dense matted deposits without any individual fibres visible (Fig 5).

**Fig 5:**
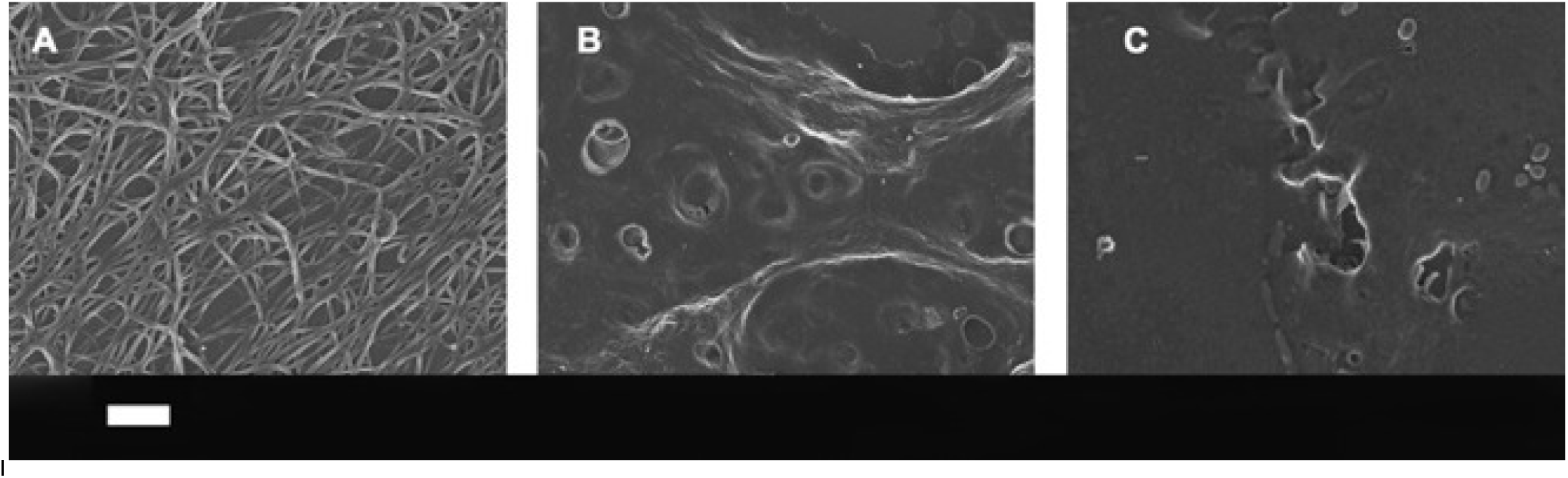
**A)** Purified fibrinogen with added thrombin but no LPS; **B)** purified fibrinogen with added O111:B4 LPS (30 seconds exposure) and 0.2 ng.L^−1^ LPS; **C)** as B but 10 minutes exposure. Scale bar: 1 *μ*m.

It is also worth rehearsing what 0.2 ng.L^−1^ of LPS means in terms of the bacterial equivalents. Watson and colleagues [46] showed in laboratory cultures that LPS amounted to some 50 fg cell^−1^ in a logarithmic growth phase, falling to 29 fg cell^−1^ in stationary phase. As remarked previously [5] this shows at once that LPS contents per cell can be quite variable, and that bacteria can shed a considerable amount of LPS at no major harm to themselves. On the basis that 1 mg dry weight of bacteria is about 10^9^ cells, each cell is about 1 pg, so 50 fg LPS per cell equates to about 5% of its dry weight, a reasonable and self-consistent figure for approximate calculations. We shall take the ‘starved’ value of 30 fg.cell^−1^. Thus 0.2 ng.L^−1^ (200 pg.L^−1^) LPS is equivalent to the LPS content of ~7.10^3^ cells.L^−1^. Most estimates of the dormant blood microbiome (summarised in [5–7]) (some are much greater [47]) imply values of 10^3^-10^4^.mL, i.e. ~1000 times smaller. In other words, a bacterial cell need lose only a small amount of its LPS to affect blood clotting in the way we describe here.

### Isothermal Titration Calorimetry

ITC is a sensitive and convenient method for detecting biomolecular interactions by measuring the heat that is released or absorbed upon binding [48]. Measurements are conducted directly in solution, without modification of immobilization of the interacting species. We used ITC to study potential interactions between human plasma fibrinogen and LPS from E. coli O111:B4. Titration of fibrinogen into LPS resulted in strong endothermic injection heats with a clear sigmoidal saturation curve indicating a direct binding interaction (Fig 6a). Assuming molecular weights of 340 kDa for fibrinogen and 20 kDa for monomeric LPS, we determined a binding stoichiometry (n) of ~0.135. This is consistent with each fibrinogen monomer binding to a micelle formed from ~75 LPS monomers. Reverse titrations were conducted, injecting LPS into plasma concentrations of fibrinogen (3 g.L-1 = 8.8 μM). Titration of 2.5 μM LPS into fibrinogen yielded endothermic injection heats greater than those observed for titration of 2.5 μM LPS into buffer alone (Fig 6b), again clearly indicating a direct binding of LPS to fibrinogen. Each injection added 125 ng of LPS into the instrument cell, increasing the LPS concentration by ~30 nM per injection. Although we expect LPS binds fibrinogen at sub-nanomolar concentrations, interactions at these concentrations are below the detection limits of the ITC instrument.

**Fig 6:**
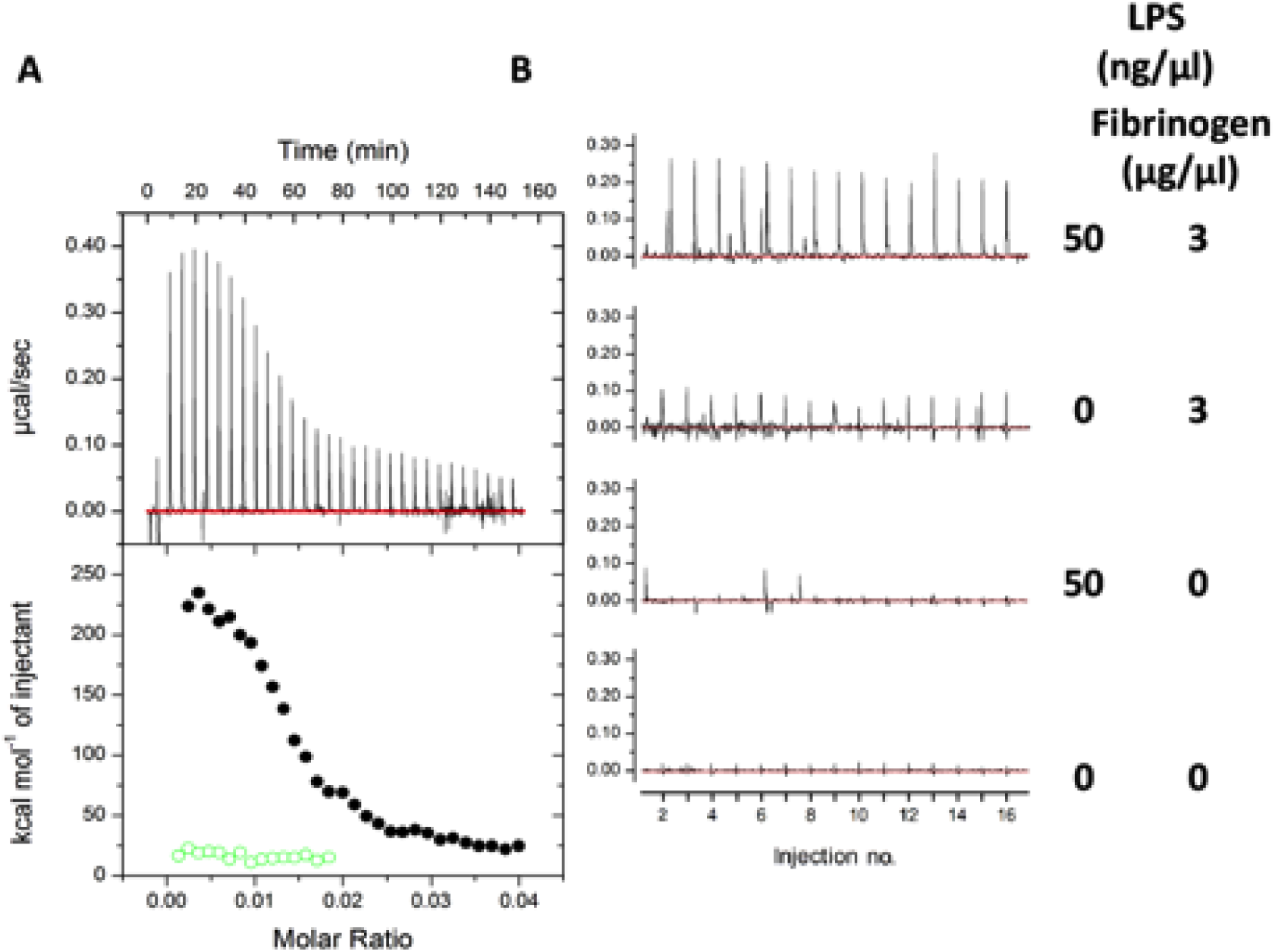
ITC analysis of the LPS-fibrinogen interaction. **A)** Titration of 8.8 μM human plasma fibrinogen *(black circles)* or buffer *(green open circles)* into 100 μM of *E. coli* O111:B4 LPS. **B)** Titration of 50 μg/μl LPS (2.5 μM) or buffer into 3 μg/μl fibrinogen (8.8 μM) or buffer as indicated. Experiments were conducted at 37 °C in phosphate buffered saline.

### Thromboelastography of whole blood and PPP

Thromboelastography (TEG) is a viscoelastic technique for measuring the clotting properties of whole blood [49, 50]. TEG of WB and PPP was performed. TEG was not performed with purified fibrinogen because the coagulation activator in the TEG is CaCl_2_ and fibrinogen is only activated by thrombin, not calcium. Fig 7 shows a typical TEG trace from a control whole blood with and without added LPS, overlaid with lines that explain the parameters extracted by the instrument and the values for those traces. The statistics are given in Table 1.

**Table 1:** Demographics of blood from healthy individuals with and without added LPS. Medians, standard deviation and p-values (values lower than 0.05 are indicated in blue) obtained using the Mann-Whitney U test are shown for iron profiles, and TEG^®^ of whole blood and plasma.

| Variables | Healthy individuals (n = 30) | Healthy individuals with added LPS (n = 30) | P-value |
| --- | --- | --- | --- |
| **AGE (years)** | 29.5 (±13.81) |  |  |
| **GENDER** |  |  |  |
| Male | 15 (50%) |  |  |
| Female | 15 (50%) |  |  |
| **IRON PROFILES** |  |  |  |
| Iron µM | 16.9 (± 6.18) |  |  |
| Transferrin g.L<sup>-1</sup> | 2.75 (± 0.46) |  |  |
| % Saturation | 25.0 (± 10.94) |  |  |
| Serum Ferritin ng.mL<sup>-1</sup> | 42.5 (± 92.47) |  |  |
| **FIBRIN FIBRE THICKNESS** | **n = 1450** | **n = 1330** |  |
| Fibre thickness (nm) | 103 (± 40) | 86 (± 82) | **< 0.0001** |
| **THROMBOELASTOGRAPHY®** |  |  |  |
| **THROMBOELASTOGRAPHY® OF WHOLE BLOOD - RECALCIFIED WITH CaCl<sub>2</sub> WITH ADDED O111:B4 LPS (10 MIN)** |  |  |  |
| MRTG (dyn.cm<sup>-2</sup>.s<sup>-1</sup>) | 2.61(± 1.13) | 2.89(± 0.90) | 0.33 |
| TMRTG (min) | 13.9 (±3.53) | 9.6 (± 3.01) | **<0.0001** |
| TTG (dyn.cm<sup>-2</sup>) | 615.0 (± 179.55) | 527.9 (± 146.65) | **0.049** |
| R (min) | 8.0 (± 1.64) | 6.2 (± 1.77) | **<0.0001** |
| K (min) | 4.9 (± 2.63) | 4.2 (± 1.23) | 0.07 |
| Angle (in degrees) | 49.8 (± 5.27) | 56.2 (± 7.02) | **0.0066** |
| MA (mm) | 55.0 (± 8.07) | 51.3 (± 6.90) | 0.092 |
| **THROMBOELASTOGRAPHY® OF PLATELET POOR PLASMA - RECALCIFIED WITH CaCl<sub>2</sub> WITH ADDED O111:B4 LPS (10 MIN)** |  |  |  |
| MRTG (dyn.cm<sup>-2</sup>.s<sup>-1</sup>) | 3.6 (± 4.35) | 4.2 (± 2.13) | 0.36 |
| TMRTG (min) | 10.6 (± 3.22) | 9.3 (± 3.68) | 0.50 |
| TTG (dyn.cm<sup>-2</sup>) | 203.9 (± 137.51) | 211.6 (± 103.67) | 0.70 |
| R (min) | 8.2 (± 2.64) | 7.1 (± 2.70) | **0.026** |
| K (min) | 4.4 (± 3.51) | 3.8 (± 2.42) | 0.18 |
| Angle (in degrees) | 63.2 (± 2.70) | 54.4 (± 10.67) | 0.23 |
| MA (mm) | 28.4 (± 8.34) | 30.1 (± 9.27) | 0.196 |
| **THROMBOELASTOGRAPHY® OF PLATELET POOR PLASMA - RECALCIFIED WITH CaCl<sub>2</sub> WITH ADDED O111:B4 LPS (30S)** |  |  |  |
|  | **n=5** | **n=5** |  |
| MRTG (dn.cm<sup>-2</sup>.s<sup>-1</sup>) | 5.94 (± 1.8) | 8.2 (± 2) | 0.166 |
| TMRTG (min) | 11.58 (± 1.2) | 9 (± 1.3) | **0.0159** |
| TTG (dyn.cm<sup>-2</sup>) | 244.4 (± 69.9) | 290.2 (± 66.5) | > 0.99 |
| R (min) | 9.8 (± 1.2) | 7 (± 1.5) | **0.031** |
| K (min) | 2.8 (± 1.6) | 2 (± 0.9) | 0.119 |
| Angle (in degrees) | 63.6 (± 6.5) | 68.8 (± 8.2) | 0.095 |
| MA (mm) | 32.8 (± 6.5) | 36.7 (± 5.5) | > 0.99 |
| **THROMBOELASTOGRAPHY® OF PLATELET POOR PLASMA - RECALCIFIED WITH CaCl<sub>2</sub> WITH ADDED O26:B6 LPS (30S)** |  |  |  |
|  | **n=5** | **n=5** |  |
| MRTG (Dynes.cm<sup>-2</sup>.s<sup>-1</sup>) | 6.3 (± 2.6) | 6.2 (± 3.6) | 0.70 |
| TMRTG (min) | 11.6 (± 2.1) | 8.9(± 1.5) | 0.15 |
| TTG (Dynes.cm<sup>-2</sup>) | 276.7 (± 43.1) | 230.8 (± 202.2) | 0.69 |
| R (min) | 9.8 (± 1.5) | 6.4 (± 1.4) | **0.05** |
| K (min) | 2.1 (± 0.5) | 2.1 (± 1.1) | 0.88 |
| Angle (in degrees) | 64.4 (± 4.3) | 70.8 (± 10.4) | > 0.99 |
| MA (mm) | 35.5 (± 3.3) | 31.5 (± 13) | 0.60 |

**Fig 7:**
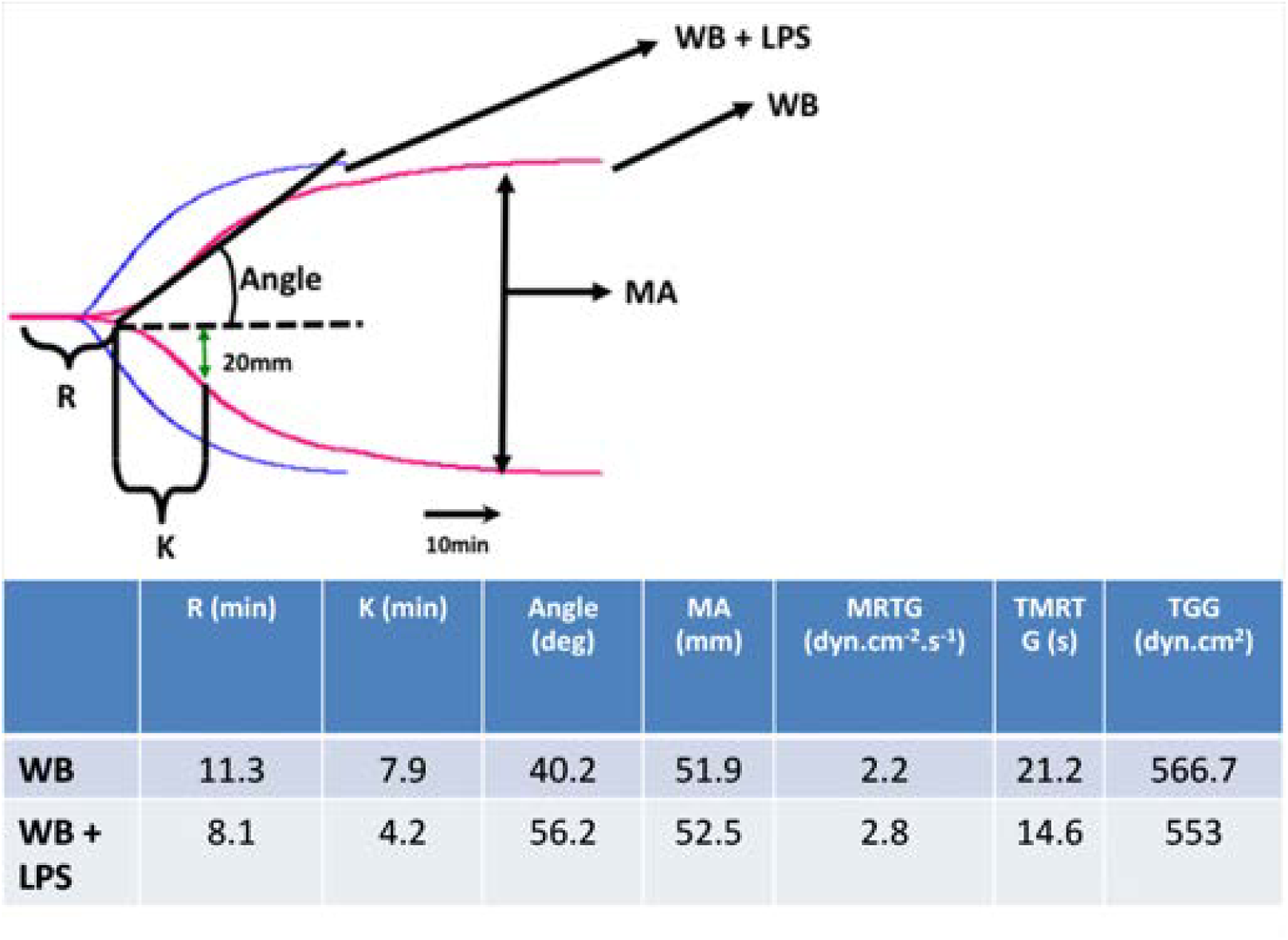
TEG overlay from a control whole blood sample with and without added LPS. R: Reaction time, first measurable clot formation; K: Achievement of clot firmness; Angle: Kinetics of clot development; MA: Maximum clot strength; MRTG: Maximum rate of thrombus generation; TMRTG: Time to maximum rate of thrombus generation; TGG: Final clot strength.

### Whole blood

TEG® analysis of whole blood (10 min incubation time with O111:B4 LPS) showed that the R, TMRTG and TGG are all significantly increased (Table 1). This suggests a hypercoagulable state as well as a denser clot, and is consistent with the SEM fibrin fibre thickness results. We have previously shown that the same concentration of LPS as used in the current paper, when added to naïve uncitrated healthy blood but <u>without added CaCl</u>_<u>2</u>_, also had an effect on coagulation after only 3 min incubation time [51]. Both TMRTG and R of naïve whole blood were also significantly shorter, also showing hypercoagulation.

### Platelet poor plasma

We also performed similar TEG experiments with PPP (see Table 1). After 10 min exposure just the initial clotting time R was changed. The results were not specific for O111:B4 LPS as O26:B6 LPS added to PPP behaved similarly. The decreased R time is indicative of reaction time, and therefore the time to first measurable clot formation is also significantly decreased (as the initiation of the clot starts faster with than without LPS), confirming a hypercoagulable state with added LPS. After 30s exposure time, both the R and the TMRTG were shorter (in the 5 patients tested). This confirms our hypothesis that LPS causes (near) instant hypercoagulability. Here also, the results were not specific for O111:B4 LPS as O26:B6 LPS added to PPP also showed a decreased R-time.

Table 1 shows a summary of the various viscoelastic properties (TEG experiments) after LPS has been added to WB and PPP. There are very substantial changes in a number of the clotting parameters. Following these 10-min exposure experiments, we shortened our experimental time to 30s and we repeated the experiments with 5 samples, using PPP, where significant changes were still observed. The results were not specific for O111:B4 LPS as O26:B6 LPS. We note that WB with added LPS showed a more pronounced change in relevant viscoelastic parameters than when LPS was added to PPP (see Table 1).

## DISCUSSION

In the introduction, we suggested that LPS might contribute to excessive blood clotting (or an activated coagulation state) via two possible routes: (i) via a direct and acute binding to plasma proteins (e.g. fibrinogen) or (ii) by an indirect or chronic (longer-term) process where it participates in an inflammatory activation via cytokine production. Here we showed that the first process is indeed possible, using tiny amounts of LPS that amounted in molar terms to less than 10^−8^ relative to fibrinogen, and demonstrated it by both viscoelastic and ultrastructural methods. We also confirmed that LPS can change the viscoelastic properties of PPP within 30s of its addition. Furthermore, WB with added LPS, but without thrombin activation, showed spontaneously formed, amyloid-like matted deposits. Purified fibrinogen experiments with O111:B4 LPS and O26:B6 LPS, with and without added thrombin showed a changed ultrastructure, suggesting that LPS indeed binds to the 340 kDa fibrinogen molecule and that the effects of this are visible ultrastructurally.

LPS, and especially its lipid A component, is highly lipophilic, and it therefore may be able to bind directly to plasma proteins, in an acute way. This might be one reason underlying the hypercoagulability [5], as well as a denser clot structure [52, 53], as seen in various inflammatory diseases. Although we show here that exposure to even tiny amounts of LPS leads to an immediate (acute) change in the coagulability parameters, we recognize that this may happen simultaneously with the chronic (longer-term) reactions (Fig 1). Fibrinogen molecules are roughly 5x45nm, and their self-assembly is a remarkable process (some 5800 are involved in generating a fibre of 80-90nm diameter and 1*μ*m length). This would explain why the highly substoichiometric binding of LPS can have such considerable effects, especially as observed in whole blood. Following Anfinsen [54] it is assumed that most proteins adopt their conformation of lowest free energy. However, this is not true for amyloid fibre formation [55] nor in the case of the autocatalytic conversion of prion protein conformations [56, 57]. At present, the exact mechanisms of action of these small amounts of LPS are not known, although it is indeed simplest to recognize fibrinogen polymerization as a cascade effect, much as occurs for amyloid and prion proteins whose initial conformation is not in fact that of their lowest free energy. Specifically [58] the ‘normal’ conformational macrostate of such proteins is not in fact that of the lowest free energy, and its transition to the energetically more favourable ‘rogue’ state is thermodynamically favourable but under kinetic control, normally (in terms of transition state theory) with a very high energy barrier ΔG^†^ of maybe 36-38 kcal.mol^−1^ [58]. Indeed, it is now known that quite a number of proteins of a given sequence can exist in at least two highly distinct conformations [59]. Typically the normal (‘benign’) form, as produced initially within the cell, will have a significant α-helical content, but the abnormal (‘rogue’) form, often in the form of an insoluble amyloid, will have a massively increased amount of β-sheet [60], whether parallel or antiparallel. In the case of blood clotting we at least know that this is initiated by the thrombin-catalyzed loss of fibrinopeptides from fibrinogen monomers (e.g. [40, 61]). The massive adoption of a beta-sheet conformation, as revealed here for the first time by the thioflavin T staining, demonstrates directly that virtually every fibrinogen molecule in the fibrin fibril must have changed its conformation hugely; it is not just a question of static ‘knobs and holes’ as usually depicted.

Previously we coined the term “atopobiotic” microbes to describe microbes that appear in places other than where they should be, e.g. in the blood, forming a blood microbiome [6]. Here we suggest that the metabolic and cell membrane products of these atopobiotic microbes correlate with, and may contribute to, the dynamics of a variety of inflammatory diseases [62–65], and that LPS, in addition to (possibly low-grade) long-term inflammation via cytokine production, may lead an acute and direct hypercoagulatory effect by binding to plasma proteins, especially fibrinogen. Specifically, we showed here that, even with very low levels and highly substoichiometric amounts of LPS, a greatly changed fibrin fibre structure is observed. An urgent task now is to uncover the mechanism(s) of this acute and immediate effect, with its remarkable molecular amplification.

## MATERIALS AND METHODS

### Sample population

30 healthy individuals were included in the study. Exclusion criteria were known inflammatory conditions such as asthma, human immunodeficiency virus (HIV) or tuberculosis, and risk factors associated with metabolic syndrome, smoking, and if female, being on contraceptive or hormone replacement treatment. Ethical clearance was obtained from the Human Ethics Committee of the University of Pretoria. Full iron tests were performed, as high serum ferritin and low transferrin levels are acute phase inflammatory protein markers [66] and indicative of inflammation. We included controls only if their iron levels were within normal ranges. Whole blood of the participants was obtained in citrate tubes and either whole blood or platelet poor plasma was used in the current study for TEG®, confocal and SEM experiments.

### LPS types, purified fibrinogen and thrombin concentration used

The LPS used was from *E. coli* O111:B4 (Sigma, L2630) and also *E. coli* O26:B6 (Sigma L2762). A final LPS concentration of 0.2 ng.L^−1^ (well below its critical micelle concentration [67] was used in all experiments bar as noted for some of the ITC measurements. It was made by vortexing a micellar suspension of 10 mg.L^−1^, followed by multiple serial dilutions. The South African National Blood Service (SANBS) supplied human thrombin, which was at a stock concentration of 20 U/ml and was made up in a PBS containing 0.2% human serum albumin. In experiments with added thrombin, 5μL of thrombin was added to 10 μL of PPP or fibrinogen. Human fibrinogen was purchased from Sigma (F3879-250MG). A working solution of 0.166 mg.mL^−1^ purified fibrinogen was prepared. This concentration was found to be the optimal concentration to form fibrin fibres in the presence of thrombin, similar to that of platelet rich plasma fibres from healthy individuals [68].

### Addition of LPS ± thrombin to whole blood, plasma and purified fibrinogen

LPS-incubated WB and purified fibrinogen was prepared for SEM without added thrombin. LPS-incubated PPP and purified fibrinogen samples were prepared as above, but with added thrombin to create an extensive fibrin fibre network.

### Isothermal Titration Calorimetry

*E. coli* O111:B4 lipopolysaccharide and human plasma fibrinogen were purchased from Sigma-Aldrich. Samples were reconstituted in warm phosphate buffered saline and incubated for 1 hr at 37 °C with shaking. LPS was then sonicated for 1 hr at 60 °C. Fibrinogen solutions were passed through a 0.2 μm polyethersulfone syringe filter and concentrations were determined by UV absorbance (E_1%_ = 15.1 at 280 nm). Samples were then diluted with buffer to the required concentration and degassed. ITC experiments were performed at 37 °C on a MicroCal Auto-iTC200 system (GE Healthcare) in high-gain mode at a reference power of 10 μcal.s^−1^, with an initial 0.5 μL (1 s) injection followed by fifteen 2.5 μL (5 s) injections with 300 s spacing. For longer titrations, the syringe was refilled and injections continued into the same cell sample. Control runs were performed in which cell samples were titrated with buffer and syringe samples were titrated into buffer, and data from these runs were subtracted from the experimental data as appropriate. Data analysis was performed in Origin, using the supplied software (MicroCal).

### Thromboelastography®

TEG® was used to study the viscoelastic properties of the participants’ blood, before and after addition of LPS. Whole blood TEG® was performed on day of collection (after 10 min incubation time with LPS) and PPP was stored in 500 μL aliquots in a -70 °C freezer. The thawed citrated PPP (340 μL) (with and without LPS – where LPS was added to PPP. Incubation time was 10 min, as with WB. Standard TEG procedures were followed with addition of CaCl_2_ to activate the coagulation process as previously described [49, 50, 69, 70]. TEG® was also performed on 5 PPP samples 30s after adding O111:B4 LPS or O26:B6 LPS.

### Confocal microscopy

Thioflavin T (ThT) was added at a final concentration of 5 μM to 100 μL PPP. A second sample was also prepared by adding a final concentration of 0.2 ng/L LPS before the addition of ThT. After an incubation time of 10 min, at room temperature and protected from light, 10 μL of the LPS-incubated PPP (with and without LPS) was mixed with 5 μL thrombin (as above) and viewed under a Zeiss LSM 510 META confocal microscope with a Plan-Apochromat 63x/1.4 Oil DIC objective, excitation was at 488 nm, and emission measured at 505-550.

### Statistical analysis

The non-parametric Mann-Whitney U test was performed using the STATSDIRECT software.

## Abbreviations

LPS - Bacterial lipopolysaccharide; TEG - thromboelastography

## ACKNOWLEDGMENTS

We thank the Biotechnology and Biological Sciences Research Council (grant BB/L025752/1) as well as the National Research Foundation (NRF) of South Africa for supporting this collaboration. This is also a contribution from the Manchester Centre for Synthetic Biology of Fine and Speciality Chemicals (SYNBIOCHEM) (BBSRC grant BB/M017702/1).

